# Early body weight gain in TALLYHO/JngJ mice predicts adult diabetic phenotype, mimicking childhood obesity

**DOI:** 10.1101/2024.01.29.577705

**Authors:** Sieglinde Hastreiter, Sandra Hoffmann, Kerstin Richter, Martin Irmler, Raffaele Gerlini, Helmut Fuchs, Valérie Gailus-Durner, Antje Körner, Martin Hrabé de Angelis, Johannes Beckers

## Abstract

Childhood obesity and type 2 diabetes are two emerging health issues worldwide. To analyze their underlying causes and develop prevention strategies, mouse models are urgently needed. We present novel insights into the polygenic TALLYHO/JngJ mouse model for diabetes. By precisely analyzing our original phenotypic data, we discovered that body weight at weaning age is the main predictor of the adult phenotype in TALLYHO/JngJ mice. The higher the weaning weight of male mice, the more likely they are to develop diabetes later in life. In contrast, a low weaning weight protected against the development of the diabetic phenotype in adults. In females, we found that high weaning body weights led to a constant higher body weight throughout life. We also showed that specifically the suckling period, rather than the *in utero* period, is crucial for the development of the metabolic phenotype in later life. We observed an earlier onset of diabetes when the mice had higher body weights at weaning, aligning with metabolic histories observed in humans. Therefore, we recommend TALLYHO/JngJ mice as a model to investigate childhood obesity and to develop prevention strategies.

**Highlights:**

- The polygenic TALLYHO/JngJ mouse model is used to investigate type 2 diabetes, but the penetrance of the phenotype is highly variable.
- We deeply analyzed our phenotype data and find that body weight at the age of weaning (BWW) is the main predictor for the obese and diabetic phenotype in TALLYHO/JngJ male mice later in life.
- We suggest that TALLYHO/JngJ male mice are an excellent and urgently needed model to study childhood obesity.
- Our data help the relevant scientific community to better control the penetrance of the diabetic phenotype in male TALLYHO/JngJ mice.

## Introduction

Childhood obesity has become an escalating health concern worldwide, affecting over 100 million children, with some countries experiencing a prevalence as high as 5% in 2015. Pediatric obesity exerts its impact across almost all organ systems, leading to long-term consequences, including cardiovascular diseases, adiposity and diabetes in adulthood (1, 2). Alongside high birth weight, another significant risk factor is rapid body weight gain during early life stages (3). Recent studies have shown that accelerated body weight gain within the first six months of life correlates positively with overweight or obesity in pre-school years (age 2-5) and pre-adolescence (age 10-12) (4).

Beyond merely identifying risk factors and associated phenotypes, understanding the underlying mechanisms that link childhood obesity to future metabolic health issues is crucial for the development of effective intervention and prevention strategies. In this context, the utilization of mouse models becomes imperative. Through our analysis of the TALLYHO/JngJ (*TallyHo*) mouse model, we demonstrate its suitability for investigating the correlation between the risk factor of ‘rapid early body weight gain’ and the escalating health concern of childhood obesity, along with its consequential metabolic effects during adulthood. Suitable models such as *TallyHo* will play a pivotal role in advancing biomedical research aimed at devising effective prevention strategies.

*TallyHo* mice represent a polygenic mouse model for type 2 diabetes (T2D), initially described in 2001, characterized by obesity resulting from an increased mass of fat pads (5). Food restriction has been found to protect *TallyHo* mice from weight gain, and energy balance data suggest that augmented body weight in male *TallyHo* mice stems from heightened food intake due to central leptin resistance, rather than from low energy expenditure (6). Additionally, hyperlipidemia has been detected in this model, along with other notable features like hyperinsulinemia and hypertrophied, degranulated pancreatic islets. Certain phenotypic traits, such as hyperglycemia (5), are restricted to male *TallyHo* mice and are typically first observed at an age of ten weeks (7). Similarly, male-limited glucose intolerance emerges at eight weeks of age and progressively worsens over time. Furthermore, adult *TallyHo* male mice develop insulin resistance (7), closely resembling several aspects of human T2D (8).

Whole genome sequence analysis revealed 98,899 SNPs and 163,720 indels unique to the *TallyHo* strain compared to 28 other inbred mouse strains (9). To identify the major loci responsible for the *TallyHo* phenotype, outcrosses of *TallyHo* mice were conducted with the mouse strains C57BL/6J and CAST/Ei, followed by backcrossing to *TallyHo* mice (B6 cross and CAST cross, respectively) (5). Genome-wide scans led to the identification of significant linkages with hyperglycemia on Chromosome 19 and 13 for the B6 cross (5, 8), which were named *Tanidd1* (*TallyHo*-associated non-insulin dependent diabetes mellitus (=NIDDM)) and *Tanidd2*, respectively. Mice homozygous for *TallyHo* alleles at both loci exhibited higher plasma glucose levels than heterozygous mice. Additionally, two loci on Chromosomes 18 and 16 were found to interact with *Tanidd1* and *Tanidd2*, thereby increasing the predisposition for elevated plasma glucose levels (5). In the CAST cross, the strongest linkage to hyperglycemia was found at other loci on Chromosome 16, designated as *Tanidd3*, along with a locus on Chromosome 19. Given a substantial overlap of the Chromosome 19 locus between the B6 cross and the CAST cross, it was suggested that it corresponds to *Tannid1*, making *Tannid1* a major locus responsible for the hyperglycemic trait in *TallyHo* mice (5).

Furthermore, in both outcross strategies, a positive correlation between body weight and blood plasma glucose levels was observed. In the B6 cross, four loci contribute to body weight, with the locus on Chromosome 19 falling within the confidence interval for *Tanidd1*, supporting the hypothesis that *Tanidd1* plays a role in hyperglycemia and body weight (5). However, it is important to note that the complex *TallyHo* metabolic phenotype likely involves many additional genetic changes.

In addition to the observation of incomplete penetrance in hyperglycemia among *TallyHo* males (5, 10), discrepancies exist regarding the precise metabolic traits observed in different laboratories. While some studies reported hyperinsulinemia and hypertriglyceridemia in four-week-old *TallyHo* mice (7), others did not find these associations (11). Moreover, data from the latter study suggested a direct influence of hyperleptinemia on pancreatic insulin secretion in *TallyHo* mice (11). Interestingly, a previous study did not observe alterations in insulin secretion during glucose tolerance tests in young *TallyHo* mice despite hyperleptinemia (7). Variations in body weight were also noted between different cohorts and both sexes of *TallyHo* mice(6, 7). Despite *TallyHo* mice being originally generated via selective inbreeding and subsequent intercross and backcross breeding, and housing the mice under standardized and specified-pathogen free (SPF) housing conditions in our facility, similar observations were made in our own *TallyHo* cohorts. Specifically, we noted highly variable degrees of penetrance for hyperglycemia, glucose intolerance, and obesity. Consequently, we sought to investigate whether environmental conditions during early life or epigenetic factors across generations could account for the varying penetrance of the metabolic phenotype in male and female *TallyHo* mice. As such, we specifically designed the subsequent study to analyze parameters of glucose homeostasis and environmental factors of individual *TallyHo* mice, rather than relying solely on group averages. This individualized approach allowed us to identify *TallyHo* mice as an excellent model for childhood obesity and subsequent metabolic health issues. This model is suitable to provide valuable insights into the underlying mechanisms of metabolic disorders that may originate from early body weight gain.

## Results

### Variable hyperglycemia in *TallyHo* male mice

To examine the variability of glycemic levels in *TallyHo* mice, we recorded blood glucose levels (BG) following a 4 to 6 hours fasting period during the first half of the light period. BG were measured weekly or bi-weekly from the age of 4 to 18 weeks. As expected, based on published observations (5, 7), female mice were generally normoglycemic with mean BG at around 120mg/dl from 4 to 18 weeks of age (Fig. 1A, pink graph). In contrast, mean BG of male mice started at around 170mg/dl at 4 weeks of age and steadily increased to approximately 280mg/dl at the age 12 weeks. Mean BG of males stayed at this level until 18 weeks of age (Fig. 1A, black graph). In particular, we noticed that males had individual fasting BG ranging from approximately 120mg/dl to almost 600mg/dl (Fig. 1B). In contrast individual BG of females were in a much more narrow range around normoglycemic levels (Fig. 1C). Next, we traced glycemic values of all individual males over time in order to understand whether these values strongly vary between time points of individual mice or whether male mice fall into distinct groups of mice with either hyper- or normoglycemia. To illustrate this in Fig. 1B, all data from mice with BG>450mg/dl at the age of 18 weeks are shown in green (severe hyperglycemic group, 23%), <250mg/dl at 18 weeks are shown in black (low glycemic group, 56%), and those with BG between 250-450mg/dl at 18 weeks are shown in blue (intermediate hyperglycemic group, 21%). Several observations can be made in this analysis: First, at 4 weeks of age BG do not yet predict the later allocation to one of the three groups. Instead, data points at this age form an intermixed cloud from all three groups. Second, males that start to develop elevated fasting BG (approximately >200mg/dl) at 6 weeks of age are determined to become members of the severe or intermediate phenotype group at 18 weeks of age. Finally, the high variability in male BG is not due to changes between measuring time points. Instead, once BG start to increase for an individual mouse, this sets the path for the development of a more severe glycemic phenotype later in life (Fig. 1B).

**Figure 1:**
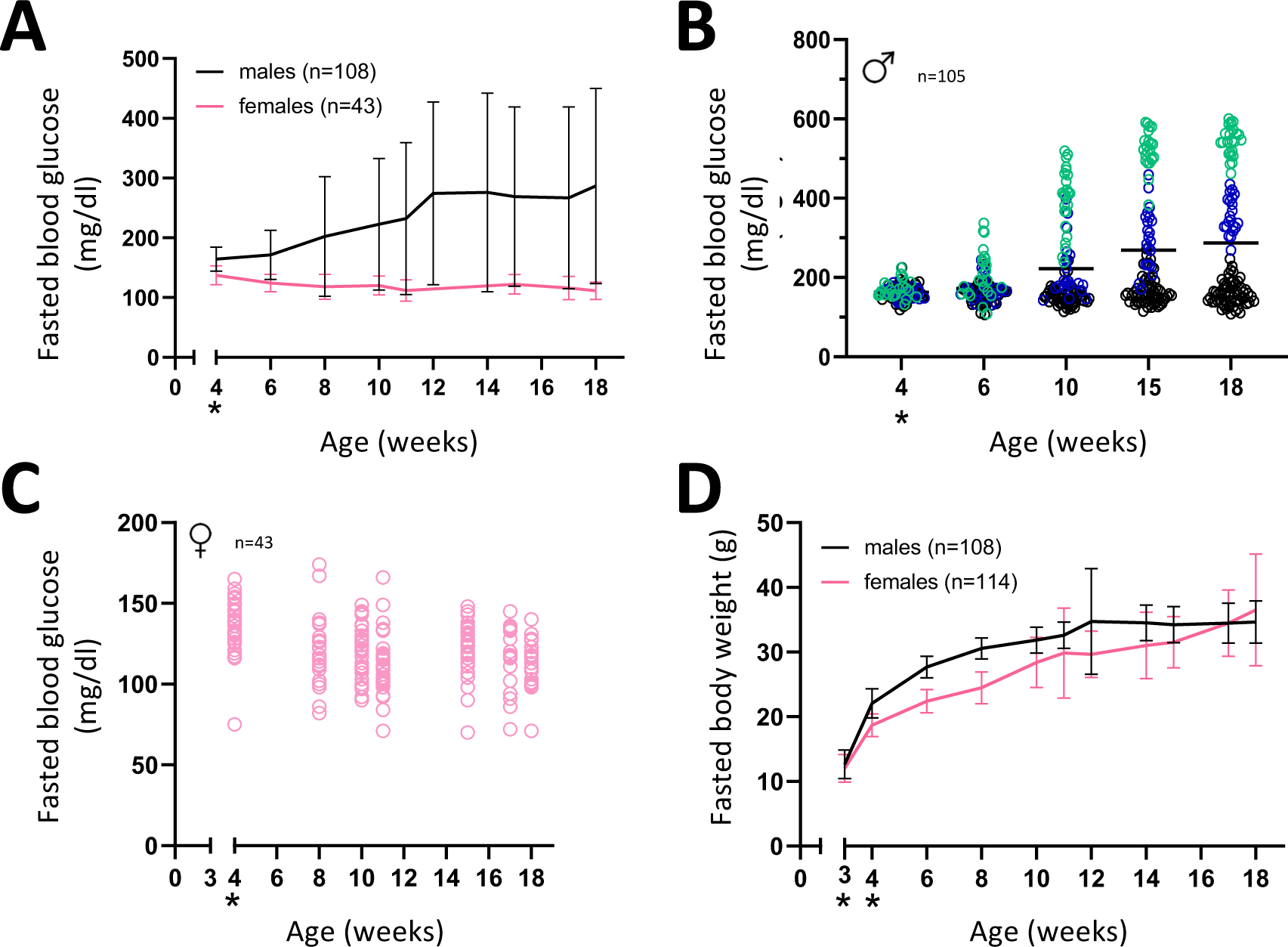
BG and BW measurements of fasted male and female *TallyHo* mice until the age of 18 weeks. Animals were not fasted (indicated *) until older than 4 weeks. BG (mg/dl) and BW (g) were measured after 4-6 hours fasting during the first half of the light period (resting phase). **(A)** Fasting BG were measured at weekly or bi-weekly intervals in males (black line) and female (pink line) *TallyHo* mice. Graphs show mean values +/- standard deviation. **(B)** Fasting BG of individual male *TallyHo* mice at ages 4, 6, 10, 15, and 18 weeks. All data points from mice that had BG above 450 mg/dl at 18 weeks are green, between 250 to 450 mg/dl at 18 weeks are blue, and below 250 mg/dl at 18 weeks are black. Horizontal bars indicate mean values. **(C)** Fasting BG of individual female *TallyHo* mice (pink) at ages 4, 8, 10, 11, 15, 17 and 18 weeks. Horizontal bars indicate mean values. **(D)** BW of female (pink) and male (black) *TallyHo* mice measured weekly or bi-weekly. Graphs show mean values +/- standard deviation. The n=108 male mice were derived from 18 litters and the n=114 female mice were born in 20 litters.

As another basic phenotypic parameter, we measured body weights (BW) from the age of weaning (3 weeks of age) until the age of 18 weeks in weekly or bi-weekly intervals (Fig. 1D). In females we observed a rather constant increase up to a mean BW of approximately 35g. Males reach this mean BW approximately at the age of 12 weeks. From this age on the mean BW of males no longer increases, presumably due to the progression to an overt diabetic phenotype in the severe hyperglycemic group (Fig. 1D).

Overall, these data confirm the incomplete penetrance of the diabetic phenotype in *TallyHo* males. However, once fasting BG start to increase between 6 and 10 weeks in individual males this is predictive of severe hyperglycemia later (see green data points in Fig. 1B).

### Litter size correlates with mating age and body weight at weaning

With the aim to search for a potential origin of the male *TallyHo* hyperglycemic phenotype we investigated earlier ages and examined litter sizes from parents of different ages and respective body weight at weaning (BWW). We found the highest mean litter sizes when parents were young (approximately 7 to 13 weeks of age at mating). Mean litter sizes became smaller when parental mice mated at ages between 14 and 19 weeks (Fig. 2A). We observed an even stronger inverse correlation between sizes of litters and BW of pups at the time of weaning. Whereas mean BWW of pups from small litters with 6 or 8 siblings were around 13g, these decreased to approximately 10g for litters with 11-15 siblings (Fig. 2B with correlation coefficient r=-0.73). This difference in BW is dramatically visible in the two mice shown in Fig. 2C, which were born on the same day, have the same father and whose mothers were sisters. Yet the smaller mouse in Fig. 2C was born and raised in a large litter and the larger mouse was born and raised in a small litter.

**Figure 2:**
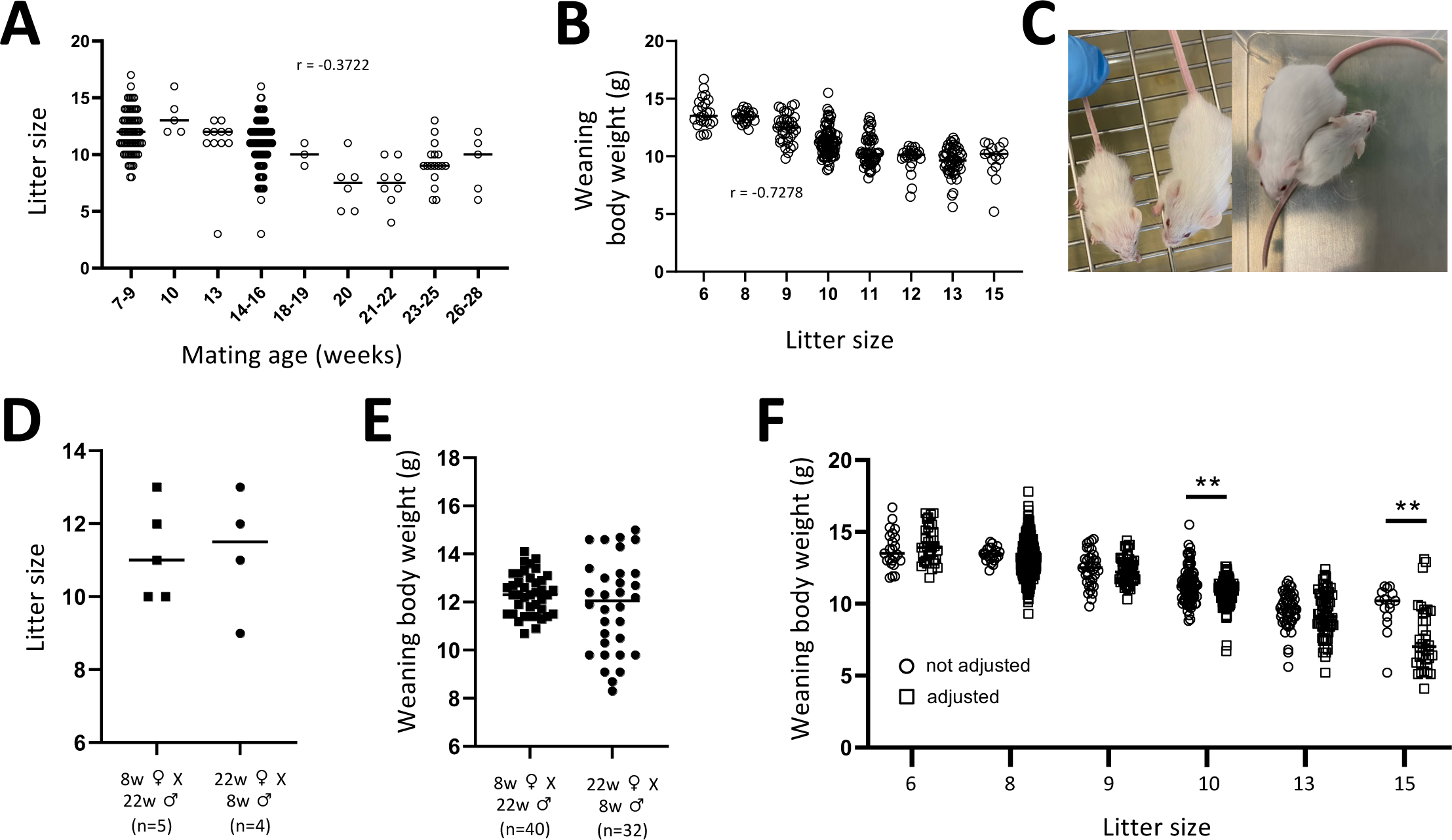
Correlations between mating age, litter size and BWW age in the *TallyHo* mouse strain. **(A)** Shows litter sizes in number of pups born dependent on the age of parents at the time of mating. **(B)** Shows BW measured at weaning age (20 – 22 days after birth) depending on litter sizes. **(C)** Shows a pair of male *TallyHo* mice born on the same date from the same father but in either a large litter (small mouse) or a small litter (bigger mouse). **(D and E)** Show litter sizes and respective BW of pups at weaning age when parents had mixed ages. As indicated in the panels, mothers were 8 weeks old and fathers were 22 weeks old at the time of mating and vice versa. Differences in both panels did not reach significance (t-test). n in **(D)** refers to number of litters, whereas n in **(E)** refers to individual mice. **(F)** Shows BW of *TallyHo* mice at the age of weaning dependent on litter sizes. Here we compare natural litter sizes (circles, not adjusted) and adjusted litter sizes (squares, adjusted). Litter sizes were adjusted either by shifting pups from their mothers to respective aunts or simply by reducing litter sizes 2 days after birth at the latest. ** indicate p < 0.01 in the pairwise t-test comparison. In all panels horizontal bars in clouds of data points indicate mean values and r is the correlation coefficient calculated with Spearman’s correlation. X-axes in these graphs do not represent scales but groups. All data were collected from male and female pups.

Although we somewhat expected that litter sizes might be more affected by the age of the mother than by that of the father, this assumption was not supported by our data. We mated 8 weeks old females with 22 weeks old males, and vice versa, and could not detect a significant difference in litter sizes between both groups (Fig. 2D). Accordingly, we also did not observe a significant difference in the mean BWW from pups of the two groups (Fig. 2E). If anything, then there might be a trend towards higher variability in BWW when mothers were older than fathers as compared to the group born from young mothers and older fathers.

We also addressed the question whether the inverse correlation between litter size and BWW has its roots *in utero* or after birth. We did this by comparing BWW between natural litter sizes versus experimentally adjusted litter sizes. Adjustments of litter sizes were done either by transferring pups from their mothers preferentially to respective aunts (that were mated with the same male and gave birth simultaneously to smaller litters) within the first 2 days after birth. However, in this analysis there is no general trend towards lower or higher BWW when adjusted litter sizes are compared to natural litter sizes. This suggests that the major impact of litter sizes on BWW occurs mostly after birth during the time of suckling and not so much *in utero* (Fig. 2F).

Overall, in *TallyHo* mice litter size decreases with the increasing mating age of the respective parents and the number of siblings negatively correlates with BWW. The effect of the litter size originates mostly during the suckling period and depends on the age of both parental mice.

### BWW as predictor of the diabetic phenotype in male *TallyHo* mice

Next, we examined whether the differences in BWW in male *TallyHo* mice corelated with the progression of fasted BG and, more generally, the progression of the diabetic phenotype. For this, animals were classified in groups according to their BWW <10g (17% of male mice), between 10-12g (33%), between 12-14g (36%), and >14g (14%). Whereas the group of mice with BWW <10g did not increase their mean fasted BG from 4 to 18 weeks of age, males with higher BWW became hyperglycemic: at the age of 18 weeks the mice with BWW of 12-14g had a mean BG level of approximately 200mg/dl, the group with 12-14g of BWW developed a mean fasting BG level of around 250mg/dl, whereas those with more than 14g had mean BG of almost 400mg/dl (Fig. 3A). Also, the distribution of individual fasted BG increased from low to high BWW. This is particularly evident at the age of 18 weeks (Fig. 3B). Especially, it appears that the group of mice with BWW >14g essentially split into two groups: a group of animals with severe hyperglycemia between 300-600mg/dl and a group with lower glycemic values between approximately 100-300mg/dl. Fig. 3C illustrates the increasing penetrance of severe hyperglycemia with increasing BWW. Whereas males with BWW <10G never developed fasted BG>400mg/dl at 18 weeks of age, approximately 40% of males developed such severe hyperglycemia. The BW curves suggest that males with weaning body weights above 12g also develop an overtly diabetic phenotype. Females with higher BWW continue to gain more weight during life as those with lower BWW (Fig. 3E). In contrast, males with weaning body weights above 12g show an attenuated gain of BW starting around the age of 10 weeks (Fig. 3D) as is characteristic for overt diabetes.

**Figure 3:**
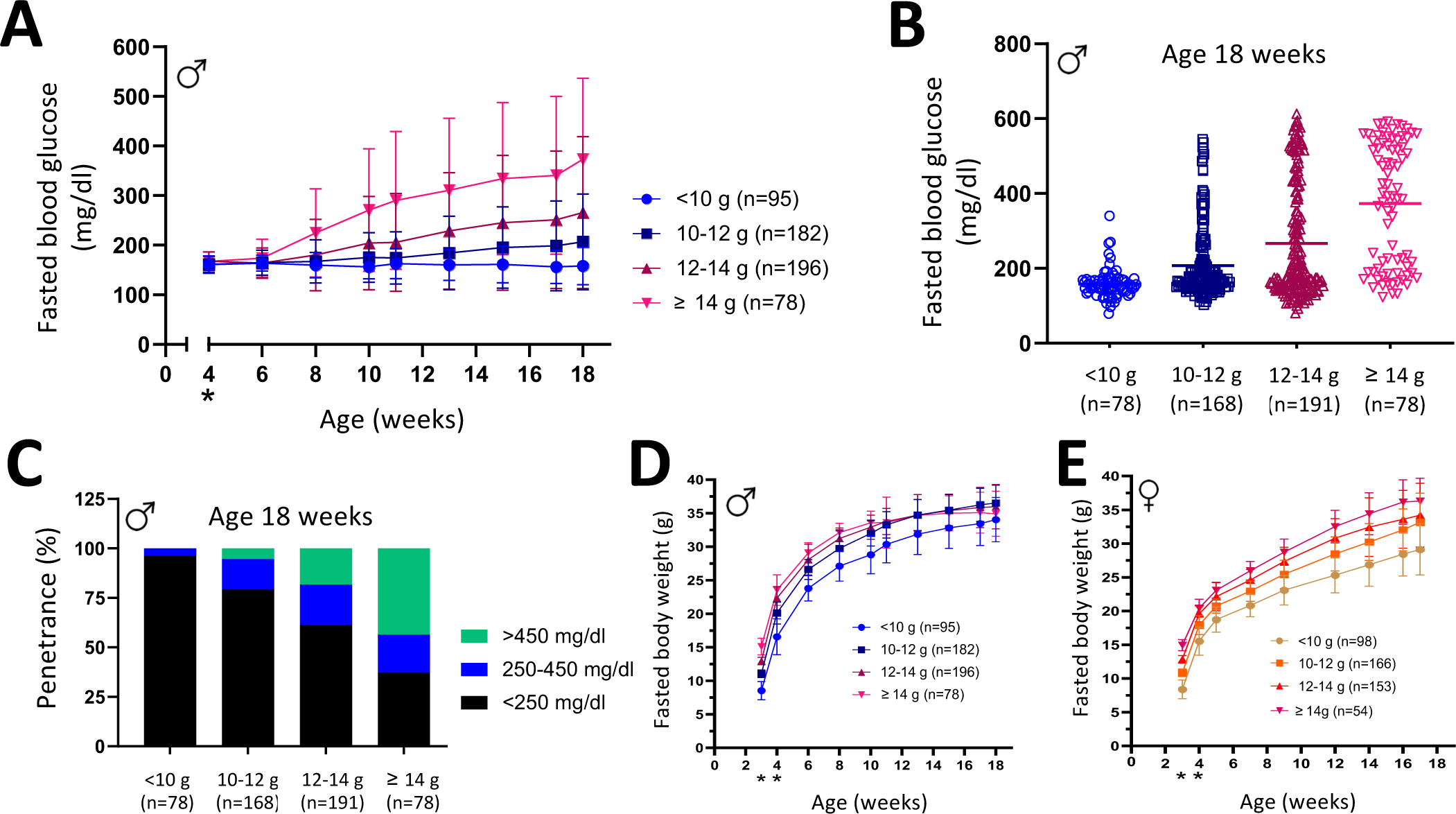
Development of fasted BG and BW of *TallyHo* mice depending on BWW (3 weeks of age). **(A)** Shows results of fasted BG measurements taken bi-weekly or weekly in male *TallyHo* mice from the age of 4 to 18 weeks. Animals were categorized according to their BWW (3 weeks) in four groups: below 10 g, between 10 g – 12 g, between 12 g – 14 g, and above 14 g. Graphs show mean values +/- standard deviation. To illustrate their distribution, **(B)** shows individual fasted BG of male *TallyHo* mice in the four weaning weight categories at the age of 18 weeks. Horizontal bars indicate mean values. Based on these data we classified fasted BG phenotypes in mild (<250 mg/dl), intermediate (250 – 450 mg/dl) and severe (> 450 mg/dl). Panel **(C)** shows the penetrance of these phenotype classes in male *TallyHo* mice at the age of 18 weeks for the four different categories of weaning body weights. Panels **(D)** and **(E)** show the BW developments of male (D) and female (E) *TallyHo* mice in the four categories of weaning body weights. Animals were not fasted (indicated *) until older than 4 weeks.

Overall, these data show that high BWW predict the development of severe hyperglycemia in *TallyHo* males and increased BW in females later in life.

### Male subgroups differ in metabolic health

To further analyze the metabolic health of *TallyHo* mice, we performed intraperitoneal glucose tolerance tests (GTT) at the age of 15 weeks and insulin tolerance test (ITT) at the age of 17 weeks. For *TallyHo* males data are analyzed for fasted BG below <250mg/dl, between 250-450mg/dl and >450mg/dl at the start of the tolerance tests (Figs. 4A and B). Even *TallyHo* males with low fasted BG show increased glycemic excursions in comparison to *TallyHo* females and C57BL/6N males (Fig. 4A). Although these mice started at rather similar fasted BG (<200mg/dl on average), their mean BG rose to about 500mg/dl at 30-60 minutes after glucose injection and did not return to initial fasted BG at the end of the GTT as in the control groups (female *TallyHo* and male C57BL/6N). Glycemic excursions of *TallyHo* males with intermediate initial fasted BG (250-450mg/dl) are even more extended and reach peaks >600mg/dl mean values after 15-60 minutes. At the end of the GTT, mean BG did not return to the fasted blood levels at the beginning of the test. Instead, the mean BG level of this group of mice after two hours was at approximately 500mg/dl (Fig. 4A). The mean BG level of *TallyHo* males with severe hyperglycemia reached peak levels after 15-30 minutes with almost 700mg/dl, whereas in the second phase of the GTT glycemic values of these severely hyperglycemic *TallyHo* mice were similar to those of the intermediate group of mice (Fig. 4A). Thus, all *TallyHo* males including those with mild fasting hyperglycemia did not reach normoglycemic levels after two hours and were glucose intolerant.

**Figure 4:**
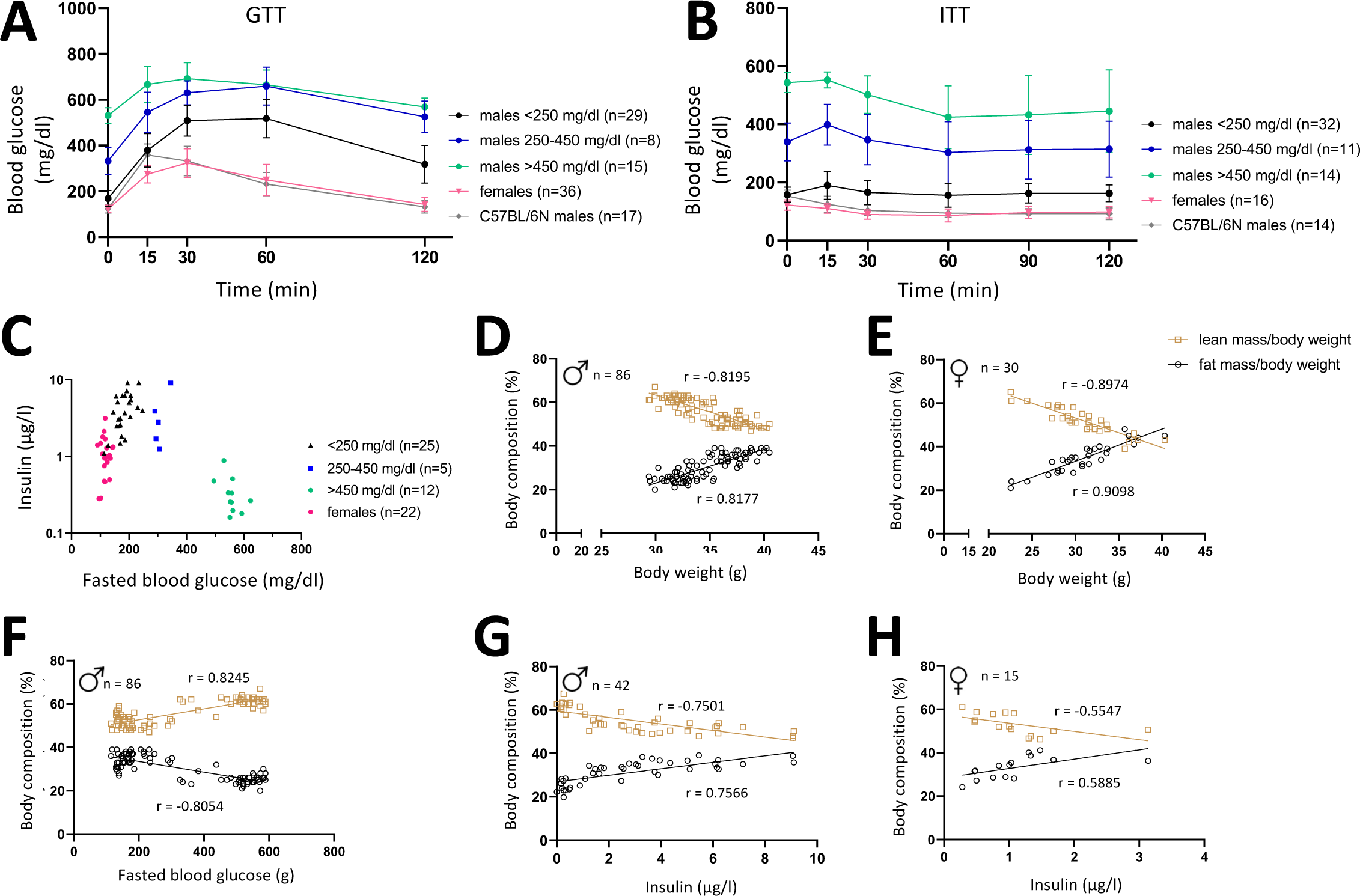
Glucose and insulin tolerance tests and body composition in *TallyHo* mice. Glucose tolerance (GTT) at the age of 15 weeks **(A)** and insulin tolerance (ITT) at the age of 17 weeks **(B)** were measured female (red) and male *TallyHo* mice, and for male C57BL/6N (as control, grey). For male *TallyHo* mice the data are shown separately for the three phenotype classes of mild (black), intermediate (blue) and severe (green) fasting BG. In panel **(C)** blood insulin concentrations are shown in relation to fasted BG concentrations at the age of 18 weeks. Classes of mice and colors used are as in panels (A) and (B). Panels **(D)** and **(E)** show body composition (in % lean, and % fat mass) in relation to the BW for male and female *TallyHo* mice. Measurements were made using NMR at the age of 16 weeks. Relative fat and lean masses are correlated to fasted BG **(F)**, and blood insulin measurements in male **(G)** and female **(H)** *TallyHo* mice. *r* is the Pearson correlation coefficient.

In the ITT *TallyHo* males with mild and intermediate hyperglycemia (<250mg/dl and 250-450mg/dl) appear to be insulin resistant. The injection of insulin after fasting did not lead to a drop in glycemic values in these mice (Fig. 4B). Remarkably, in *TallyHo* males with severe hyperglycemia (>450mg/dl at start of ITT) the insulin injection reduced mean BG after 30 minutes and reduced them even further after 60 minutes. In *TallyHo* mice with intermediate initial BG the response to insulin injection was lower, but like the severe hyperglycemic males, they reached the lowest BG 60 minutes after the insulin injection (Fig. 4B). In the control groups (female *TallyHo* and male C57BL/6N), the insulin injection led to an immediate drop in BG until the lowest levels were reached after 60 minutes (Fig. 4B).

It appears that *TallyHo* males with severe hyperglycemia have developed an overt diabetic phenotype. Accordingly, they show low blood insulin levels even at high fasted BG (Fig. 4C), suggesting that pancreatic beta-cells might no longer produce or release sufficient insulin. In contrast to these mice, blood insulin levels of *TallyHo* males with mild or intermediate hyperglycemia had elevated blood insulin levels compared to *TallyHo* females (Fig. 4C).

Regarding the body composition measured by NMR at 16 weeks of age, we observed in both, male and female *TallyHo* mice, a positive correlation between fat mass and BW. Mice with higher BW have increased relative fat masses and reduced relative lean masses (Fig. 4D males, Fig. 4E females). Additionally, males with high fasted BG show reduced fat mass/BW in comparison to males with lower fasted BG (Fig. 4F). Accordingly, blood insulin concentrations positively correlate with the fat mass/BW in *TallyHo* males (Fig. 4G) and females (Fig. 4H).

Overall, *TallyHo* males with mild or intermediate fasting hyperglycemia are characterized as pre-diabetic with glucose intolerance, elevated insulin levels and increased fat mass. Interestingly, these mice showed no or low response to insulin injection. Whereas males with severe fasting hyperglycemia are overtly diabetic, characterized by glucose intolerance, accompanied with decreased blood insulin and reduced body fat mass. In addition, insulin injection leads to decreased BG in overtly diabetic *TallyHo* males.

### *TallyHo* food intake increases with overt diabetic phenotype

So far, our data show that the higher penetrance and severity of the diabetic phenotype in *TallyHo* males correlates with an increased BWW and lower litter sizes. Next, we wanted to understand whether this is followed by differences in food consumption later in life, in particular in the comparison between males with severe versus mild diabetic phenotypes.

For this, we analyzed food and water consumption of 6 and 18 week old males with BWW <10.5g (n=25) and >13.5g (n=22) in metabolic cages for 21 hours. Interestingly, the young males showed no significant differences in food intake when the groups with low and high BWW were compared (Fig. 5A). In particular, males that later became highly hyperglycemic (>450mg/dl, green data points in Fig. 5A) did not consume more food at the age of 6 weeks than males with high BWW that later in life had lower BG. This changed dramatically by the time these mice were 18 weeks old. At this age males that became highly hyperglycemic (>450mg/dl) consumed significantly more food than males that had BWW <10.5g or as males with comparable BWW but with lower BG (Fig. 5A).

**Figure 5:**
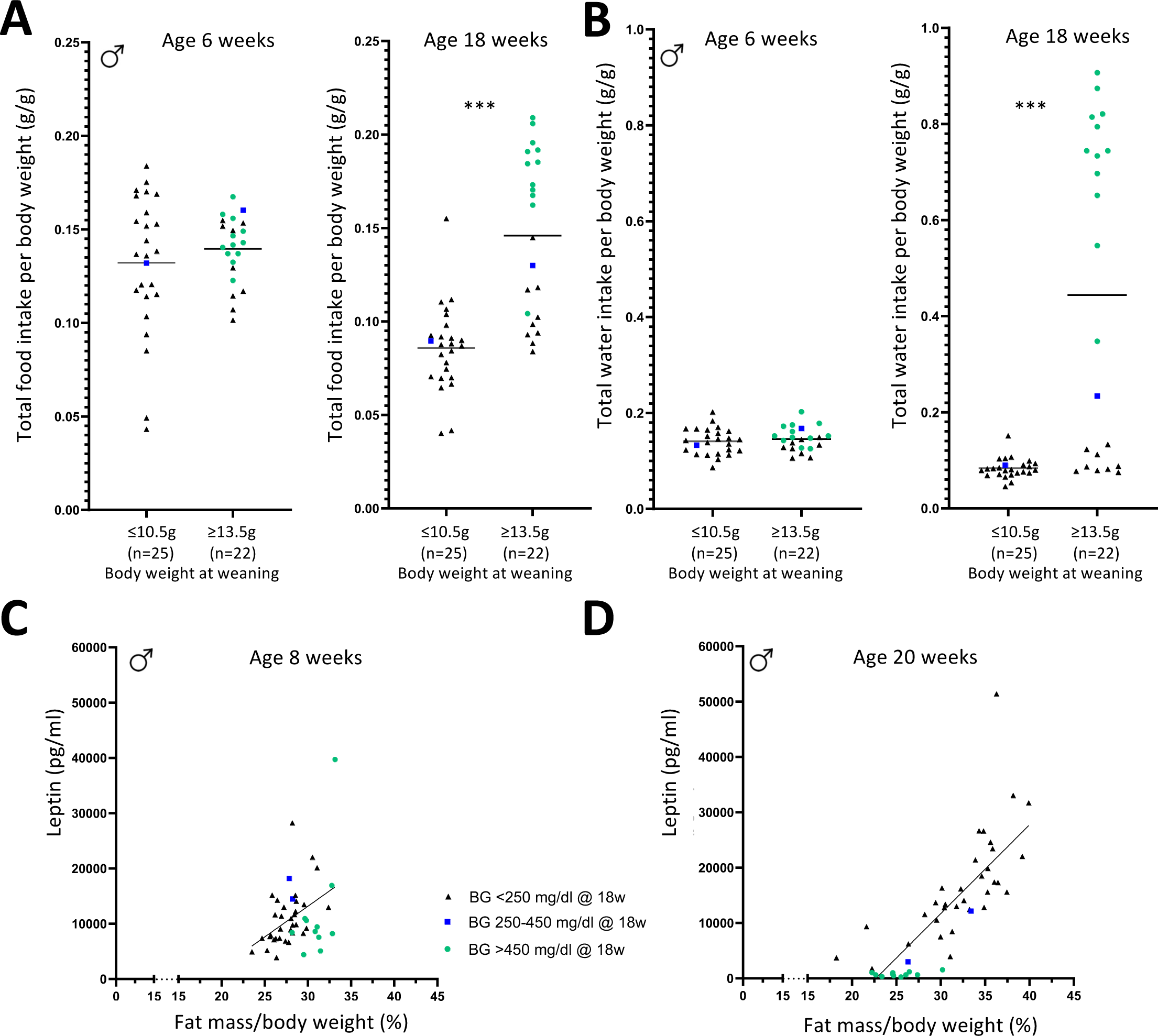
Hyperphagia and polydipsia in diabetic male *TallyHo* mice. Food **(A)** and water **(B)** consumption of male *TallyHo* mice at 6 and 18 weeks of age measured in metabolic cages. Measurement was performed on two consecutive days for each 21 hours. Data presented are mean values of both measurements. *** indicate p<0.001 in the Mann-Whitney test, horizontal lines indicate mean values. Blood leptin concentrations in male *TallyHo* mice at 8 weeks **(C)** and 20 weeks **(D)** of age. Line indicates linear regression; *r* is the Pearson correlation coefficient (8 weeks: *r* = 0.3910, 20 weeks: *r* = 0.7867).

This was paralleled with similar differences with regards to water consumption. Water consumption was highly significantly increased only in males with BWW >13.5g and that developed severe hyperglycemia by the age of 18 weeks and not at the age of 6 weeks (Fig. 5B). The occurrence of polydipsia in hyperglycemic, lean, insulin deficient male *TallyHo* is an additional indication of an overt diabetic phenotype in these mice. No such differences in food and water consumptions were observed in females (not shown).

As expected from their lean phenotype, highly hyperglycemic males at the age of 20 weeks were highly deficient in blood leptin levels. In contrast, at the age 8 weeks these same mice still had blood leptin levels that were in the same range as for *TallyHo* males that later developed no or a mild diabetic phenotype (Fig. 5C and D). No such losses of leptin signaling in the blood were observed in females (not shown). Also, we did not observe any differences in leptin sensitivity tests depending on the severity of the diabetic phenotype in males at the age of 19 weeks (data not shown).

Taken together, we observe that juvenile male *TallyHo* mice (6 weeks) that later develop overt diabetes consume food and water at normal quantities. Only later in life (18 weeks) when males have developed overt diabetes, they become hyperphagic and polydipsic.

### Altered gene expression between pre-diabetic and diabetic TallyHo mice

Next, we analyzed gene expression variations underlying the spectrum from pre-diabetic to overt diabetic phenotypes each in mice with >13,5g BWW. This analysis is provided as Supplementary Material and includes Suppl. Fig. 1. In summary, we observed varying degrees of differential gene expression (padj<0.1) in visceral adipose tissue (6440 genes), liver (781), muscle (67), hypothalamus (82), and subcutaneous adipose tissue (0) at 18 weeks of age.

## Discussion

In this study, we have uncovered novel insights into the phenotypic variations within the *TallyHo* mouse model. Our findings reveal that body weight at weaning serves as a key predictor for the adult phenotype, with mice raised in small litters and consequently exhibiting higher body weight at weaning being more susceptible to developing an overt diabetic phenotype in adulthood. Conversely, we observed that low body weight at weaning appears to confer protection against the development of an overt diabetic phenotype in male *TallyHo* mice. These observations shed light on potential causes for the reported differences in penetrance and phenotypic outcomes observed in various laboratories (6, 7, 11).

Considering that non-diabetic male *TallyHo* mice have limited utility when investigating this model as a representative of human T2D, our findings offer a novel approach to enhance the penetrance of the desired phenotype. This aspect is of particular interest to researchers working with *TallyHo* mice, as it opens new avenues to increase the relevance and applicability of the model for the study of human type 2 diabetes.

At the outset of this project, we embarked with the hypothesis that *TallyHo* mice could serve as a polygenic model for exploring the potential of epigenetic inter-generational inheritance of obesity and diabetes (12–15). This hypothesis was initially based on the intriguing observation that progeny of older parents, displaying a more severe metabolic phenotype, exhibited higher penetrance and a more severe phenotype, in contrast to progeny from young parents (Fig. 2 and data not shown). However, our meticulous analysis clearly demonstrates that the differences in penetrance and severity of obesity and diabetes are primarily attributable to variations in litter sizes and consequently different body weights at weaning. Therefore, our findings caution against hasty interpretations of inter-generational inheritance. As pointed out also by others, careful consideration of possible confounding factors is essential when exploring potential epigenetic inheritance across generations, and additional investigations are warranted to unravel the underlying mechanisms (16, 17).

In general, the manifestation of a severe diabetic phenotype of *TallyHo* males is characterized by hyperglycemia, low insulin levels in the blood, reduced body fat mass, and glucose intolerance. Consistent with previous reports, we also observed an increase in body weight until the onset of hyperglycemia, particularly in male *TallyHo* mice (Kim 2006). Notably, as hyperglycemia developed over time, we observed hyperphagia and polydipsia in these mice.

Despite increasing evidence that litter size profoundly impacts adult metabolic health in mice (18–20), it is not yet standard practice for studies involving mouse models to routinely report litter sizes or weaning body weight information. However, in the case of the *TallyHo* mouse model, litter size, and consequently body weight at weaning, are critical determinants for the later severity of the diabetic phenotype. In light of our findings, we strongly recommend the routine reporting of body weights at weaning for mice used in experiments, especially when examining metabolic phenotypes. Unlike reporting litter sizes as previously suggested (19), body weight at weaning is an individual-specific parameter, not a group measurement, and enables a more precise evaluation of lifetime phenotype data.

In addition to providing important information to researchers working with the *TallyHo* model, our findings consistently draw parallels to the concerning issue of early childhood obesity. Childhood obesity can profoundly impact glucose homeostasis, leading to insulin resistance and type 2 diabetes not only during adulthood but also during adolescence (1). Furthermore, longitudinal studies have revealed that a significant majority of children with obesity at the age of 3 years continue to be obese or overweight also in their adolescence (21). This bears a striking resemblance to our observation in *TallyHo* males, where higher body weight at weaning correlates with an earlier onset of hyperglycemia or, in case of constant normoglycemia, a sustained higher body weight throughout life (Fig. 3).

In addition, the *TallyHo* male mice exhibit similarities to the human situation where not all obese adults develop insulin resistance and type 2 diabetes, instead, some individuals remain metabolically healthy.

In a recently published article, the association between infancy growth rates and overweight/obesity in greek pre-school children (2 – 5 years old) and adolescents (10 – 12 years old) were analyzed. The authors demonstrated a positive association between rapid weight gain during infancy and overweight/obesity at pre-school years and adolescence, with statistical significance in the latter (4). In this study, infancy was defined as the first 6 months of life, which roughly corresponds to the suckling period in our mouse model. Although *TallyHo* mice did not display differences in birth weight (data not shown), the high body weight at weaning could be considered equivalent to the rapid weight gain during infancy in humans. Notably, our observations regarding body weight development align with the trends and effects reported by Moschonis et al., further supporting the proposition that the *TallyHo* model holds promise as a suitable mouse model for studying childhood obesity.

In addition to providing novel insights for researchers employing the *TallyHo* mouse model and the potential to enhance the penetrance of the diabetic phenotype, we believe this mouse model holds value in investigating the escalating issue of childhood obesity and its ensuing consequences, offering a means to explore strategies for preventing metabolic health issues in later life.

## Materials and Methods

*Animal welfare and approval:* All mice were housed as described: https://www.mouseclinic.de/about-gmc/mouse-husbandry/index.html. Experiments were approved by Bavarian authorities (Regierung von Oberbayern, Sachgebiet 54). All animal experiments described in this study were performed in strict accordance with the ethical guidelines of Helmholtz Munich. All animals were monitored by animal welfare officers and every effort was made to minimize suffering and reduce the number of animals used.

*Breeding:* Male *TallyHo* mice were mated for 5–7 days with one or two females. Litter size adjustment or shifting of litters (pups of the same father were exchanged between mothers to achieve small or large litters) was performed within 48 hours after birth. Animals for each experiment were bred in different time-shifted batches to analyze independent replicates.

*BW and BG:* Weekly BW measurement began at three weeks of age. Biweekly fasted BG assessment started from six weeks of age (4–6 hours fasting, starting around 8 a.m.), unfasted BG was measured at an age of four weeks. Contour BG analyzers (Bayer Vital) and corresponding sensor strips (Ascensia Diabetes Care) analyzed tail blood samples. The cohort for indirect calorimetry underwent weekly fasted BG and BW measurements from four weeks of age, with unfasted BW recorded at three weeks of age.

*Glucose tolerance test:* Mice were fasted for 4–6 hours, followed by basal BG and BW measurement. Subsequently, 2g glucose/kg BW (20% glucose solution, B. Braun Melsungen) was intraperitoneally injected, BG were recorded at 15, 30, 60 and 120 minutes post-injection. *NMR:* MiniSpec LF60 (Bruker Optics) measured lean and fat mass in unfasted mice, along with BW recordings.

*Insulin tolerance test:* Mice fasted for 4–6 hours, followed by basal BG and BW measurement. Then, 1lU/kg BW insulin ((10µl insulin (Huminsulin 100, Lilly) in 10ml 0.9% NaCl (Mini-Plasco, B. Braun Melsungen)) was intraperitoneally injected. BG were recorded 15, 30, 60, 90 and 120 minutes post-injection.

*Indirect calorimetry:* Males aged 6 and 18 weeks were single housed in a home cage indirect calorimetry system (TSE Systems) with ad libitum access to water and normal chow. Monitoring occurred for 21 hours on two consecutive days (1 p.m. to 10 a.m. the following day). Food and water intake was measured every 20 minutes. Total 21-hour food intake was normalized to BW.

*Blood samples and organ withdrawal:* Mice at 18 weeks were anesthetized using isoflurane (IsoFlo, zoetis) for retrobulbar blood samples. Plasma was obtained from the blood collected in EDTA tubes (KABE Labortechnik) after centrifugation (2,000xg, 4°C, 5 minutes). Organs were collected after cervical dislocation; hypothalamus was extracted on a cooling plate. Tissue and plasma samples were flash frozen in liquid N_2_, then stored at -80°C.

*ELISA:* Insulin ELISA (Mouse Insulin ELISA, Mercodia) and Leptin ELISA (Quantikine Mouse/Rat Leptin Immunoassay, R&D Systems) followed manufacturer’s protocols. Insulin ELISA used undiluted plasma samples, while Leptin ELISA used 1:20 dilution.

## Author contributions

SHa, SHo, KR, MI, and RG performed experiments and analyzed data. Manuscript was written by SHa and JB. KR, MI, HF, VGD, AK and MHdA revised or contributed significantly to the manuscript. JB and MHdA conceived and designed research studies.

## Acknowledgments

We thank Dr. Sibylle Sabrautzki for expertise and support regarding animal welfare issues, Ann-Elisabeth Engelniederhammer and Andreas Mayer for outstanding technical assistance and the animal care takers of Helmholtz Munich for professional assistance. This work was supported by grants from the German Center for Diabetes Research (DZD e.V.) and the German Federal Ministry of Education and Research (Infrafrontier grant 01KX1012 to MHdA). The authors declare that they have no conflict of interests. Funders had no role in data collection, interpretation, and reporting.

